# Stratified Random Sampling Methodology for Observing Community Mask Use within Indoor Settings: Results from Louisville, Kentucky during the COVID-19 Pandemic

**DOI:** 10.1101/2021.02.25.432837

**Authors:** Seyed M. Karimi, Sonali S. Salunkhe, Kelsey B. White, Bert B. Little, W. Paul McKinney, Riten Mitra, YuTing Chen, Emily R. Adkins, Julia A. Barclay, Emmanuel Ezekekwu, Caleb X. He, Dylan M. Hurst, Martha M. Popescu, Devin N Swinney, David Johnson, Rebecca Hollenbach, Sarah Moyer, Natalie C. DuPré

**Author notes:** Corresponding author: Seyed M. Karimi. These Authors had an equal contribution to the manuscript (Seyed M. Karimi, Sonali S. Salunkhe, Kelsey B. White). 1. Seyed M. Karimi. 2. Sonali S. Salunkhe. 3. Kelsey White. 4. Bert B. Little. 5. W. Paul McKinney. 6. Riten Mitra. 7. YuTing Chen. 8. Emily R. Adkins. 9. Julia A. Barclay. 10. Emmanuel Ezekekwu. 11. Caleb X. He. 12. Dylan M. Hurst. 13. Martha M. Popescu. 14. Devin N. Swinney. 15. David A. Johnson. 16. Rebecca Hollenbach. 17. Sarah Moyer. 18. Natalie C. DuPre. **Competing Interests**: The authors have declared that no competing interests exist.

## Abstract

Wearing a facial mask can limit COVID-19 transmission. Measurements of communities’ mask use behavior have mostly relied on self-report. This study’s objective was to devise a method for measuring the prevalence of mask-wearing and proper mask use in indoor public areas without relying on self-report. A stratified random sample of retail trade stores (public areas) in Louisville, Kentucky, USA, was selected and targeted for observation by trained surveyors during December 14−20, 2020. The stratification allowed for investigating mask use behavior by city district, retail trade group, and public area size. The average mask use prevalence among observed visitors of the 382 visited public areas was 96%, while the average prevalence of proper use was 86%. In 17% of the public areas, at least one unmasked visitor was among the observed visitors; in 48%, at least one improperly masked visitor was observed. The average mask use among staff was 92%, but unmasked staff were observed in fewer public areas, as an unmasked staff member was observed in 11% of the visited public areas. The average prevalence of proper make use among staff was 87%, similar to the average among visitors. However, the percentage of public areas where at least one improperly masked staff was observed was 33. Significant disparities in mask use and its proper use were observed among both visitors and staff by public area size, retail trade type, and geographical area. Observing unmasked and incorrectly masked visitors was more common in small (less than 1500 square feet) public areas than larger ones, also in food and grocery stores than other retail stores. Also, the majority of the observed unmasked persons were male and middle age adults.

## Introduction

Transmission of respiratory viral infections like COVID-19 can be reduced considerably by using a facial mask.^1-6^ Therefore, understanding mask-wearing practices in a community is important and can inform public health policies and interventions, especially with a lack of continuous random COVID-19 testing. However, common methods of measuring mask-wearing practice are suspected of significant inaccuracy, particularly surveys based on self-report,^7-11^ resulting in attenuation bias. When non-interventional observations of mask use replace self-reporting, non-representative sub-populations (e.g., university students and clinic population) have been studied.^12-14^ Observational studies that do not focus on a specific sub-population are scarce.^15^ This study’s primary aim was to develop a method to measure the prevalence of mask-wearing in public accurately. To this purpose, visitors and staff of a representative sample of indoor public areas (PAs) in the city of Louisville (estimated 2019 population: 766,757),^16^ Kentucky (USA) were observed. The representativeness of PAs was ensured by a stratified random sampling method, which allowed for assessing disparities in mask usage across city districts, industries, and PA sizes. In contrast to existing mask usage studies,^7-15^ this study also measured the prevalence of proper facial mask use.

### Materials and methods

An observational method was developed to assess facial mask-wearing practice in Louisville, Kentucky, during December 14−20, 2020. The observational aspect of this study meant that it was not experimental but relied on systematic observation of subjects’ behavior in indoor public areas without intervention.^17^ The set of observed PAs was selected using a stratified random sampling technique from the pool of retail trade businesses in the city. Surveyors were assigned subsets of the randomly selected PAs and were trained to log their time in a PA with a standardized assignment sheet and to submit observations with a standard online questionnaire (see “The questionnaire and data collection method” and “The survey implementation” sections below for details). The filled questionnaires were cross-checked with surveyors’ assignment sheets then the data was downloaded and refined to calculate four prevalence proportions of mask use: (1) the proportion of unmasked among visitors, (2) the proportion of unmasked among staff, (3) the proportion of incorrectly masked among visitors, and (4) the proportion of incorrectly masked among staff.

### Ethics committee approval

This study was funded by the Louisville Metro Department of Public Health and Wellness (LMPHW) through the Coronavirus Aid, Relief, and Economic Security Act (the CARES Act) and approved by the University of Louisville Institutional Review Board (IRB# 20.966).

### Definitions

In this study, an indoor public area was defined as an establishment (e.g., a business, store, or facility) where the public can go without an appointment and need to be assisted or attended by the establishment’s staff.

A PA’s staff were identified by their uniform, displaying the PA’s proprietary brand. If the PA’s staff did not use uniforms, location and action (e.g., working behind counters, service area desks, and checkout registers) were used to identify them.

Any type of facial covering—such as bandanna, cloth mask, neck gaiter, disposable surgical mask, cone-style mask, N95, and other respirators—used to protect against aerosol transmission of infectious particles was defined as “mask” or “facial mask.”

A person was considered “masked” if the person wore a facial mask that covered both nose and mouth, “incorrectly masked” or “improperly masked” if either nose or mouth was not covered, “unmasked” if both nose and mouth were not covered.

### Public area stratification criteria

Indoor PAs were selected using a random sampling technique stratified by the city district, industry, and PA size in order to observe mask-wearing in various parts of the community, in different types of PAs and PA sizes. Seven city districts—namely, (1) South & South West, (2) West Center, (3) North West, (4) North Center, (5) Central, (6) South East, and (7) East & North East—were constructed based on the geographical proximity of zip codes, demographic and median income(**Appendix Table 1, Appendix Fig 1**). Notably, people who are black are more highly concentrated in specific areas of the city, as the city suffers from a legacy of racial segregation highlighted discriminative zoning practices.^18-20^ According to data from the 2010 census, the non-Hispanic black population concentration was the highest in the North West district of the city (75%: 54%−93% in different zip codes). This district had the highest proportion of children (28% in 2010), and the median household income was the lowest ($22,848: $16,686−$27,565 in 2018) among all districts. The districts with the second and third largest share of non-Hispanic black population were West Center (33%) and Central (23%). The West Center and Central districts’ median household income was the city’s second and third lowest: $37,469 and $43,911 in 2018, respectively. The non-Hispanic black population in other districts was between 6% and 11% in 2010. The city’s wealthiest district was East & North East, with a $91,141 median household income (**Appendix Table 2**). The U.S. median household income in 2018 was $63,179.^21^

The sampling strategy then considered industry for the second stratification criteria. Among Standard Industrial Classifications (SIC), Retail Trade (SIC division G, codes 52xxxx−59xxxx) was considered in this study.^22^ Observing the other nine industrial classifications (e.g., Mining, Construction, Manufacturing, etc.) would have required business-specific arrangements and resulted in higher research costs than what was available to this study. Among the subcategories of Retail Trade, Automotive Dealers and Gasoline Service Stations (SIC codes 55xxxx) and Eating and Drinking Places (SIC codes 58xxxx) were excluded to preserve the observational and non-interventional nature of the study, as most businesses in these two types of retail provide personal assistance to customers. Therefore, the study focused on the remaining subcategories of Retail Trade: Building Materials, Hardware, Garden Supply & Mobile Home Dealers (SIC codes 52xxxx), General Merchandise Stores (SIC codes 53xxxx), Food Stores (SIC codes 54xxxx), Apparel and Accessory Stores (SIC codes 56xxxx), Home Furniture, Furnishings and Equipment Stores (SIC codes 57xxxx), Miscellaneous Retail (SIC codes 59xxxx). The last category included drug stores and proprietary stores, liquor stores, used merchandise stores, and books stores, among others. A surveyor could conveniently visit a typical business classified under any of the six retail industries as a customer with no purchase and time cost for the surveyor.

The third stratification criterion was PA size so that the team could investigate if mask-wearing behavior differed in larger PAs where visitors may spend more time and cluster in greater numbers than smaller PAs.

### Businesses’ data and surveying clusters

Data on information regarding the indoor PAs in the city were obtained from Data Axle in ArcGIS format.^23^ A total of 4,648 PAs in Louisville were classified as Retail Trade (excluding SIC codes 55xxxx and 59xxxx) and were considered for observation. Information on the PAs was imported into the statistical software STATA 16.0 (STATACorp, LLC, College Station, TX, USA), which included zip code, SIC code, company name, primary address of the retail store, secondary address of the retail store, the retail location’s latitude and longitude, and the square footage area of the retail store in ten categories.

Three variables representing stratification criteria were constructed: district, retail groups, and PA size. The city zip codes were grouped as described above, and the seven districts were formed (**Appendix Table 1, Appendix Fig 1**). Six retail trade classifications were regrouped into four based on the similarity of trade types. Specifically, SIC codes 52xxxx (Building Materials, Hardware, Garden Supply & Mobile Home Dealers) and 57xxxx (Home Furniture, Furnishings and Equipment Stores) were combined to form the furniture, equipment, and home improvement stores group, and SIC codes 53xxxx (General Merchandise Stores) and 56xxxx (Apparel and Accessory Stores) were combined to form general merchandise and apparel stores group. 827 PAs were grouped into furniture, equipment, and home improvement stores, 996 in general merchandise and apparel stores, 878 in food and grocery stores, and 1,947 PAs in miscellaneous retail stores.

In terms of size, PAs were categorized into four size groups: PAs with square footage (1) less than 1499, (2) between 1500 and 2499, (3) between 2500 and 4999, and (4) greater than 5000. The specific recategorization was to increase the uniformity of PAs’ distribution over size, given the footage brackets in the Data Axle dataset. In practice, PA size groups 1, 2, 3 and 4 included 1186, 1360, 954, and 1148 PAs, respectively.

Ultimately, 112 surveying clusters were formed. Each cluster included PAs specified by a combination of either of the seven districts, four retail groups, and four PA sizes (**Appendix Table 3**).

### Sample selection

The number of PAs in the 112 clusters ranged from 9 to 151. Tertiles of the distribution of the number of PAs in clusters were ≤ 25, > 25-44, and > 44. Three PAs were randomly selected without replacement from any cluster that fell in the first tertile, four PAs from the second tertile clusters, and five PAs from the third tertile clusters (**Appendix Table 4**). As a result, 447 PAs we randomly selected to be observed. Observing 447 PAs was perceived feasible based on the survey’s cost assessment and funding. Using the same method, an extra 447 PAs were selected as a back-up in case any of the indoor PAs were non-operational.

To demonstrate the selected PAs’ representativeness for observation, the proportion of each district in all selected PAs with the district’s human and PA populations proportions (**Appendix Table 5**). The differences in proportions between the selected PAs and human population were small, between −2.7% and 2.7% across the seven districts, and the differences between selected PAs and PA population proportions were also small, between −3.8% and 4.3%. This suggested that the selected PAs were representative of the population distribution and PA distribution across districts. As a more detailed representativeness test, the difference in each district’s share in all selected PAs of a specific size was calculated with the district’s human and PA populations shares. Therefore, two measures of error (equal to the absolute value of the two differences) were calculated for each of the 28 district-PA size clusters. When the shares of selected PAs in the districts were compared to the districts’ human population shares, the mean error was 1.98%, and quartiles of the errors were 1.43%, 1.95%, and 3.05%, respectively. Comparing the share of selected PAs in the districts to the districts’ PA population shares resulted in a mean error of 2.98%, with quartiles cut-points of 1.24%, 2.96%, and 4.48%.

### Surveyors’ assignment sheet

This study employed ten trained surveyors. Each surveyor observed mask use in a subset of the randomly selected 447 PAs and a subset of the 447 back-up PAs that were also randomly selected. A surveyor’s assignment sheet was a Word document that included two tables: a table of selected PAs and a table of reserve PAs. Each table had six columns: Unique ID, PA Zip Code, Retail Group, PA Size, PA Name, PA Address, Observation Number, and Notes.

A surveyor was asked to observe only the table of selected PAs. If a selected PA was not operational, then the surveyor was asked to replace it with a PA in the same retail group and the same size from the reserve table.

The Unique ID column was pre-populated with three-digit numbers. Unique IDs were generated to connect the collected data from a PA to its other information reported in the Data Axle master dataset. Surveyors would insert the PA’s unique ID into the online questionnaire; hence the unique ID will appear as a survey data column.

An observation number is another randomly generated number that a surveyor transferred from the online questionnaire to the assignment sheet. The observation number was used as an extra identification tool to connect a PA questionnaire (hence the survey data) to the PA’s notes on the assignment sheet. Notes on the assignment sheet may contain data correction remarks if necessary (e.g., if a surveyor inserted a piece of information about the PA incorrectly or a selected PA was not operational).

### The questionnaire and data collection method

The survey’s questionnaire included three sections: (1) PA Information, (2) Observing Visitors, Observing Staff, and each section included five questions. The first section included pre-answered questions on PA identification information, Date of Survey, and Time of Survey. Also, a surveyor was asked to enter the PA Name and PA Unique ID from the assignment sheet (**Table 1**).

**Table 1:**
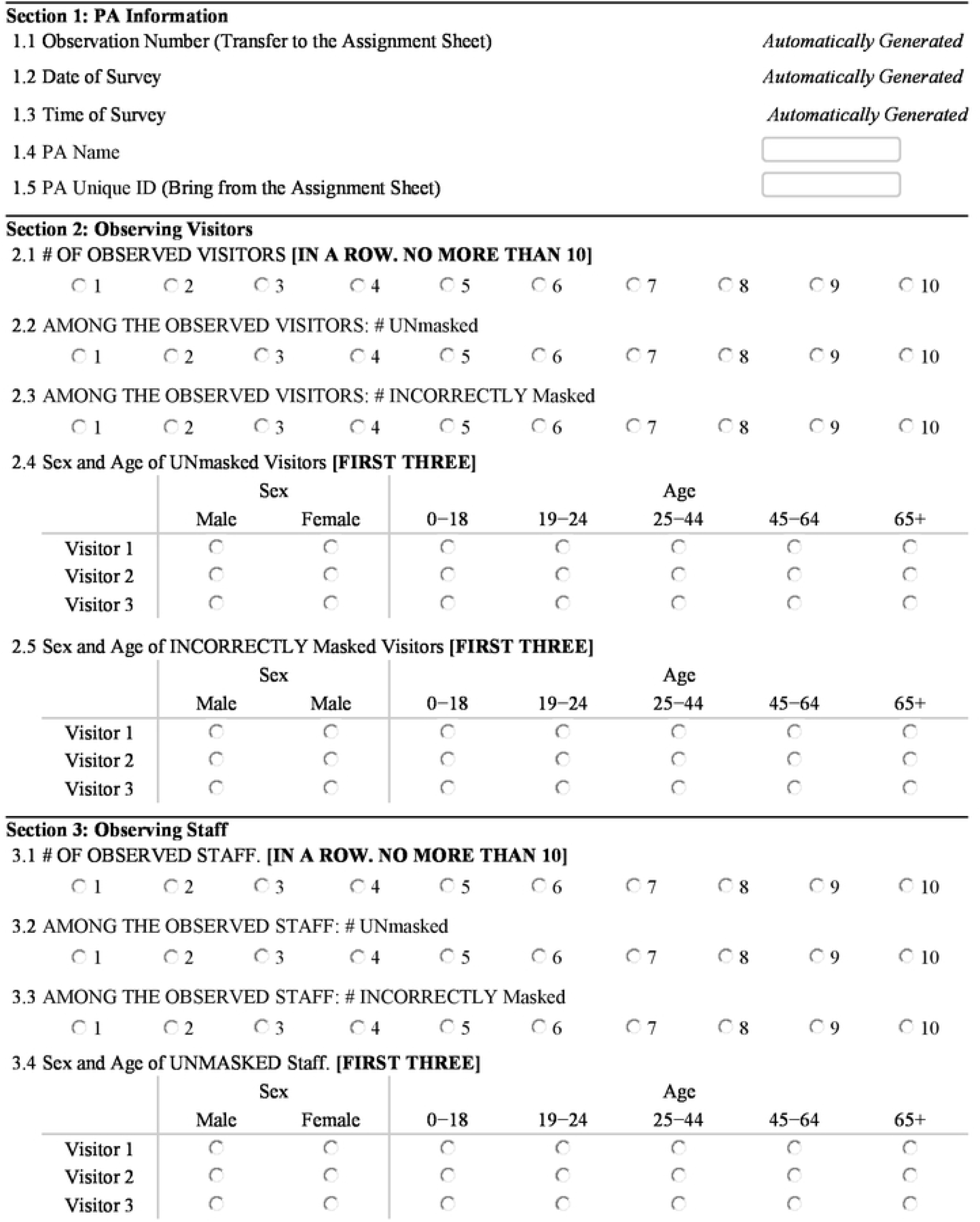
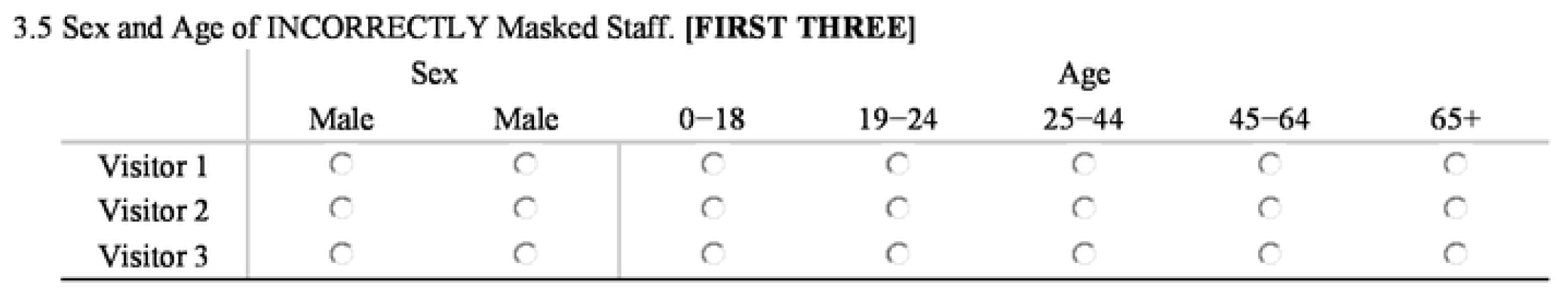
The survey questionnaire

In the second section, a surveyor was asked to observe the mask use of up to ten visitors in the PA. That is, if there were more than ten visitors, they needed to observe only ten of them, but if there were fewer than ten visitors, they needed to observe all of them. They could start observing as soon as they entered the PA or start observing at any time or location after entering the PA. In any case, they (1) should not wait to start their observation after seeing an unmasked or incorrectly masked visitor and (2) needed to observe ten visitors consecutively. The answer to this question determines the denominator of the proportion of mask users and incorrect mask users among visitors. The second and third questions asked the numbers of the unmasked and incorrectly masked among the observed visitors, respectively. The fourth and fifth questions asked the surveyor’s to perceive the sex and approximate age-ranges (0−18, 19−24, 25−44, 45−64, and 65+ years) of the unmasked and incorrectly masked. In the third section of the questionnaire, a surveyor was asked about mask use among the PA staff. The section was structured similarly to the second section (**Table 1**).

The survey’s questionnaire was designed in Qualtrics (Qualtrics, Provo, UT), and its link was emailed to surveyors to record observations from a PA. When a questionnaire was submitted, the link could be used to start filling another questionnaire for another PA. Answers were stored and were transferred to STATA for statistical analyses.

### The survey implementation

Surveyors were trained to use the Qualtrics link and complete the questionnaire. The definitions of a mask, an unmasked person, and an incorrectly masked person were reviewed. Moreover, examples of a typical observation were presented to them, and their questions about the questionnaire and the data collection process were answered before the survey began and during the survey. The survey was conducted from December 14 through December 20, 2020. A surveyor visited each indoor PA as a customer between 10:00 a.m. to 6:00 p.m. EST and observed mask-wearing behavior. Surveyors spent 5 to 15 minutes in a PA, depending on the store size, and completed the electronic survey.

## Results

During the survey week, 382 PAs were observed. The largest number of PAs were observed in the North Center district (n=71, 19%) (**Appendix Fig 2**). This district also included the largest share of the city’s population (20%) (**Appendix Table 4**). The smallest number of PAs were observed in the North West district (n=32, 8%) (**Appendix Fig 2**); the district included 11% of the city’s population (**Appendix Table 4**). The observed PAs included 19% furniture, equipment, and home improvement stores, 24% general merchandise and apparel stores, 35% food and grocery stores, and 23% miscellaneous retail stores (**Appendix Fig 3**). A majority of the surveyed locations were over 5000 square feet in size (43%) or between 2500 and 5000 square feet (21%) (**Appendix Fig 4**).

The distribution of the number of observed PAs across the study’s clusters was not precisely the same as the distribution of the number of selected PAs (**Appendix Tables 4 and 6**). In effect, the difference between a district’s share in total observed PAs and the district’s share in total human and targeted PA populations was greater than that for the district’s share in total selected PAs (**Appendix Table 7**). Specifically, the difference between the observed PAs and human population shares was between −4.2% and 5.6% across the districts, and the difference between observed PAs and PA population shares was between −4.9% and 5.6%.

The larger error in observation than selection was the result of two implementation challenges. Firstly, several smaller-sized PAs either were not operational or could not be observed without an appointment. Secondly, the number of PAs in furniture, equipment, and home improvement stores and in miscellaneous retail stores that could not be observed without an appointment was greater than such PAs in other trade groups.

### Mask wearing for public areas’ visitors

The mean proportion of unmasked observed visitors across all of the observed PAs was 4% (Standard Deviation (SD) = 14%), and in 83% of the PAs, there were no unmasked visitors. Both mean and variation of incorrect mask usage were greater than those of no mask use: 14% wore a mask incorrectly (SD = 22%). In 52% of the PAs, there were no observations of incorrectly masked visitors (**Fig 1a**). Incorrectly masked and unmasked visitors were most frequently observed within small public areas, those consisting of less than 1,500 square feet, where on average 10% (SD = 27%) were unmasked and 17% were incorrectly masked (SD = 29%) (**Fig 1a**).

**Figure 1:**
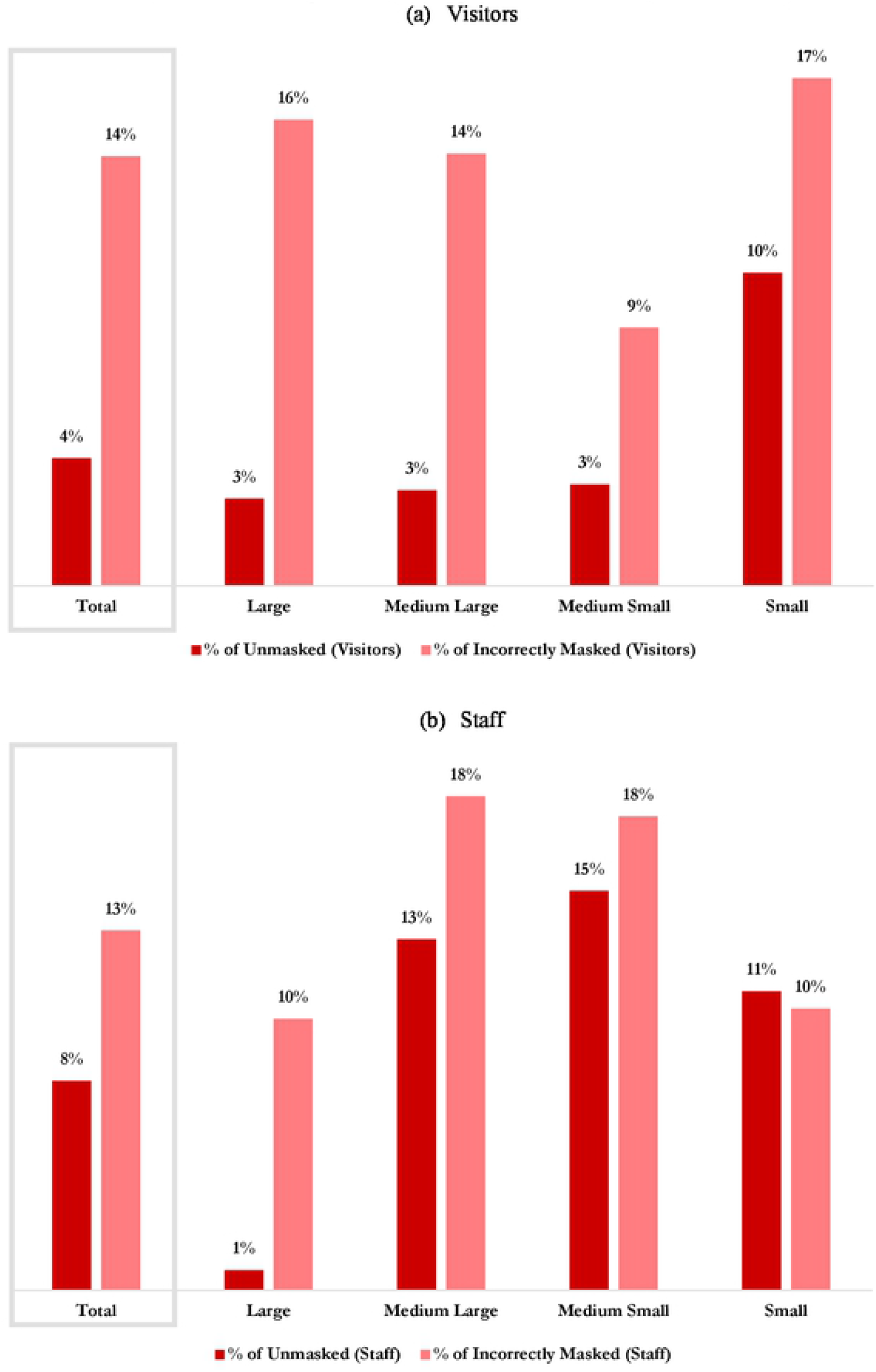
The average prevalence of the unmasked and improperly masked among visitors and staff of Louisville indoor public areas by public area size, Dec. 14-20, 2020. *Note:* **Large PAs are those with a square foolage of at least 5000, between 2500 and 4999 sq ft for medium large, 1500 to 2499 sq ft for medium small, and 1499 or less for small**.

Unmasked visitors were less frequently observed in the visited PAs from furniture, equipment, and home improvement stores, where the unmasked prevalence among visitors was zero in 92% of them, and the mean prevalence was 2% (SD=13%). In PAs from the other retail groups, the prevalence of no mask-wearing was about 4% to 5% (**Fig 2a**). The incorrectly masked visitors were most commonly observed in food and grocery stores than the other three observed retail groups. In 57% of food and grocery stores, at least one incorrectly masked visitor was among the observed visitors, and the mean proportion of incorrectly masked visitors was 18% (SD=23%) (**Fig 2a**).

**Figure 2:**
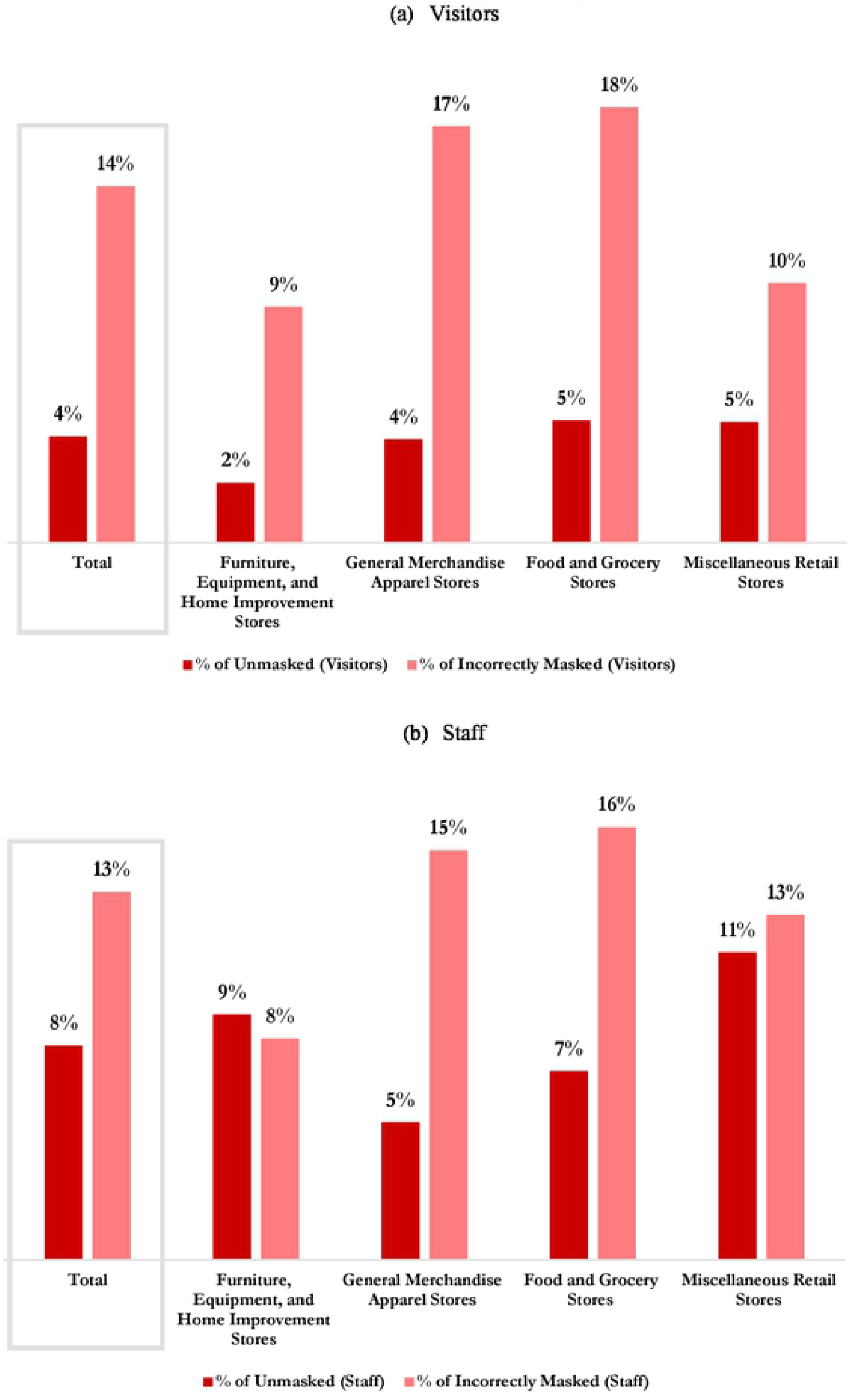
The average prevalence of the unmasked and improperly masked among visitors and staff of Louisville indoor public areas by retail trade group, Dec. 14-20, 2020

Unmasked visitors were observed most frequently in the South & South West district, where the mean prevalence of unmasked visitors was 8% (SD = 21%) (**Fig 3a**). Nonetheless, the prevalence of unmasked visitors was zero in 76% of the observed PAs in this district. Incorrectly masked visitors were observed most commonly in the North West district, where the mean prevalence of incorrectly masked visitors was 20% (SD = 29%), and at least one incorrectly masked visitor was observed in 57% of this district’s PAs (**Fig 3a**).

**Figure 3:**
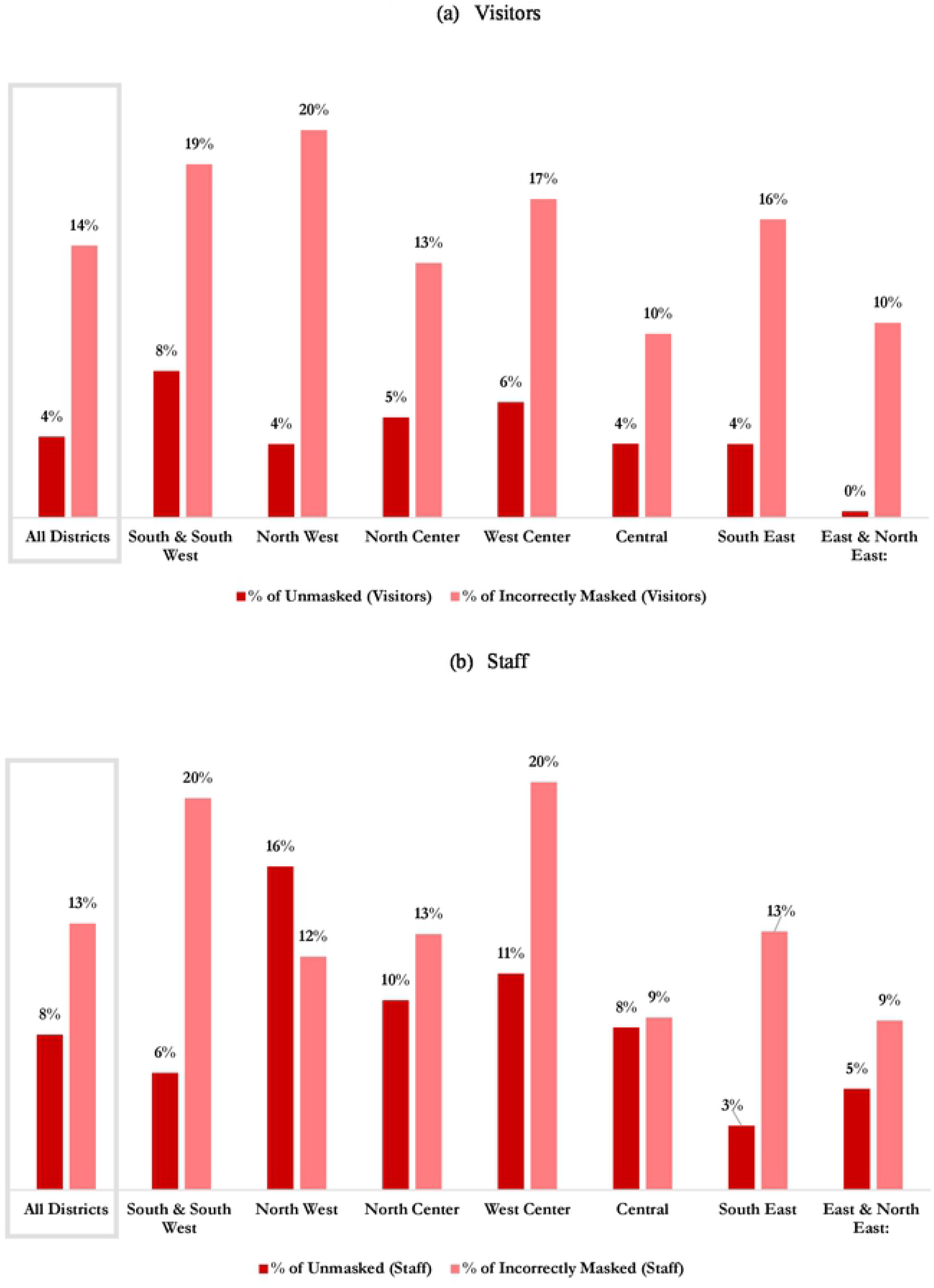
The average prevalence of the unmasked and improperly masked among visitors and staff of Louisville indoor public areas by district, Dec. 14-20, 2020

Of all observed unmasked visitors, 61% were perceived as male (**Fig 4a**). Among visitors wearing masks incorrectly, 53% were perceived as male (**Fig 4b**). About 50% of the unmasked and 48% of the incorrectly masked visitors were perceived as middle age adults (**Fig 5**).

**Figure 4:**
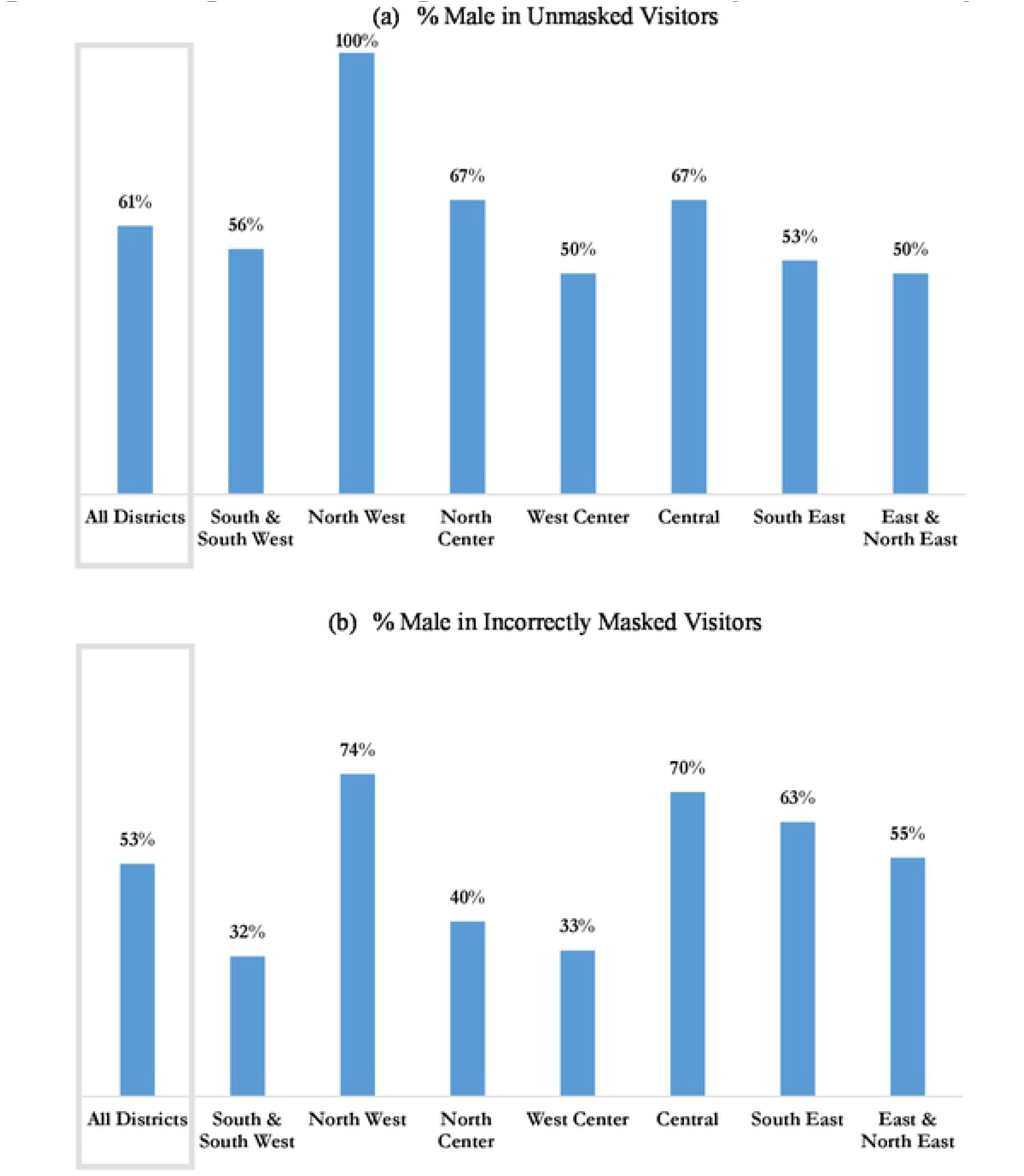
Percentage of males among unmasked and incorrectly masked visitors by district

**Figure 5:**
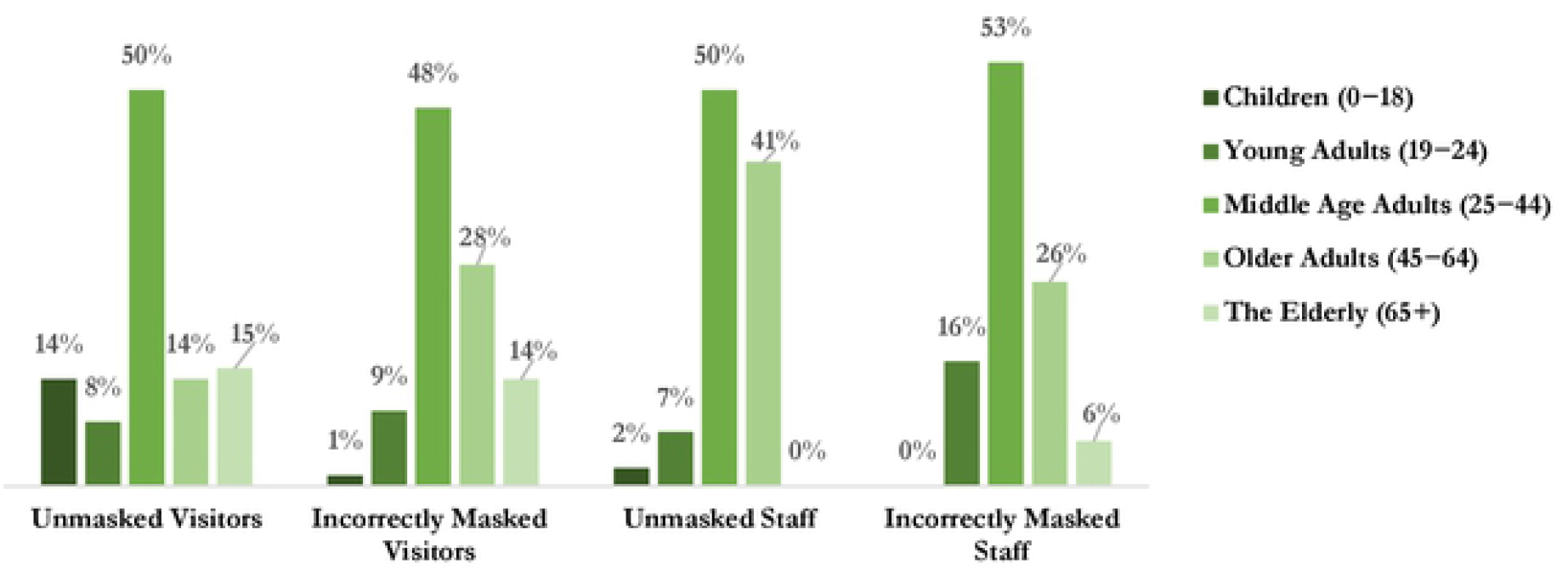
The age distribution of the unmasked and incorrectly masked among visitors and staff

### Mask wearing for public areas’ staff

Among the observed staff in the observed PAs, the mean prevalence of unmasked and incorrectly masked staff was 8% (SD=25%) and 13% (SD=25%), respectively (**Fig 1b**). The proportion of the unmasked and incorrectly masked were zero in 89% and 67% of the PAs, respectively.

Incorrectly masked and unmasked staff were most frequently observed within medium-small public areas, those with the square footage between 1500 to 2500, where the mean prevalence of no mask usage in staff was 15% (SD=33%) and prevalence of incorrect mask usage in staff was 18% (SD=30%) in medium-small PAs (**Fig 1b**).

Unmasked staff were less commonly observed in the visited PAs from general merchandise and apparel stores, where the unmasked prevalence among staff was zero in 93% of them, and the mean prevalence of unmasked staff was 5% (SD=21%). In PAs from the other retail groups, the prevalence of no mask usage was higher, between 7% and 11% (**Fig 2b**). The incorrectly masked staff were most frequently observed in food and grocery stores than the other three observed retail groups. In 36% of food and grocery stores, at least one incorrectly masked staff was among the observed staff, and the mean prevalence of incorrectly masked staff was 16% (SD=27%) (**Fig 2b**). Unmasked staff were observed most commonly in the North West district, where the mean prevalence of unmasked staff was 16% (SD=35%) (**Fig 3b**). Nonetheless, the prevalence of unmasked staff was zero in 68% of the observed PAs in this district. Incorrectly masked staff were observed most frequently in the South & South West and West Center districts-20% (SD=33%) and 20% (SD=31%), respectively (**Fig 3b**). Nonetheless, the proportion of incorrectly masked staff of these districts was zero in 65% and 59% of the observed PA in the two districts, respectively. Of all observed unmasked staff, 50% were male. Among observed incorrectly masked staff 42% were male (**Appendix Fig 5**). About 50% of the unmasked and 53% of the incorrectly masked visitors were perceived as middle age adults (**Fig 5**).

## Discussion

This study was conducted to estimate the prevalence of mask use and its proper use in indoor public areas of the city of Louisville, Kentucky (USA) in December 2020 and to explore potential patterns in mask-wearing behavior. Results suggested there was a high prevalence of mask use: 96% of visitors and 92% of staff wore a mask in the 382 observed PAs. Improper masking appeared more common than not wearing a mask at all in December 2020. Among the masked visitors and staff, respectively, 86% and 87% used it properly, covering both the nose and mouth (**Fig 1**).

While the mean mask use among staff was smaller than that among visitors (92% versus 96%), the proportion of PAs where at least one unmasked staff was observed was smaller than the share of PAs where at least one unmasked visitor was observed (11% versus 17%). Similarly, the share of PAs where improperly masked staff was observed was smaller than the share of PAs where improperly masked visitors.

This research suggests that variation in the proper use of masks may exist depending on the size of the PA (**Fig 1**). Although previous research suggests a similar finding,^24^ why this occurs remains unknown. One explanation could be the way individuals follow perceived social norms.^25^ Perhaps individuals more intentionally follow a norm in areas with more people; there is a greater expectation to follow a pre-set norm, such as wearing a mask correctly, in larger rather than smaller public areas. Others suggest that health behaviors are more specifically connected to one’s appraised threat such that the lower the appraised threat, the less likely the individual will adhere to protective behaviors.^26,27^ In other words, if one assesses a lower risk of contracting COVID-19 infection in a smaller establishment due to the location’s smaller capacity, then they are less motivated to completely adhere to those protective behaviors.^26^ Future studies examining mask-wearing behavior should include data points about individual behavior beliefs, protective beliefs, beliefs about infectious diseases, and misinformation beliefs.^28^

The findings showed that unmasked visitors and incorrectly masked visitors ranged from 2% to 5% and 9% to 18%, respectively, across different retail stores (**Fig 2**). Also, unmasked and incorrectly masked staff in the retail stores ranged from 5% to 11% and 8% to 16%, respectively (**Fig 2**). A survey conducted by the Pew Research Center in August 2020 revealed that approximately 85% of people reported wearing a mask over the past month, which has increased from 65% seen in June.^8^ The odds of observing a person wearing a mask in an urban or suburban retail store were four times than those seen in the rural areas.^29^ Despite mask mandates in place, some people either do not wear a facial mask or wear them incorrectly, thus putting not only others in their close proximity but also themselves at risk of contracting the infection due to the possibility of viral transmission from asymptomatic COVID-19 carriers. ^29^

Significant geographical variation in mask use and proper use prevalence was observed in this study. The mean unmasked prevalence varied from 0% to 8% in visitors and 3% to 16% in staff. Geographical variation in mask use was strongly correlated with income such that the districts with highest prevalence of unmasked and improperly masked individuals were among the most economically disadvantaged districts of the city. For example, the highest visitor improperly masked prevalence (20%) and the highest staff unmasked prevalence (16%) were recorded in the city’s poorest district, the North West district, where the median family income was $22,848 in 2018. Conversely, the lowest visitor unmasked and improperly masked prevalence (0% and 10%, respectively) and the lowest staff unmasked prevalence (5%) were recorded in the richest district of the city, East & North East where the median family income was $91,141 in 2018 (Appendix Table 2). Populations with low-income often face greater challenges obtaining resources to practice healthy behaviors and often face complex social contexts^30,31^, and having the financial means to purchase or make an appropriately fitted mask is likely to be an obstacle in lower income communities. Researchers have reported a higher prevalence of high-risk health behaviors among communities with low-income.^32^ In other words, although smoking and physical inactivity differ from mask-wearing, the communities facing the greatest health challenges for systemic reasons are also struggling with masking. Further, researchers currently suggest that case and mortality rates are elevated for populations with complex health challenges, in areas health disparities already exist, and potentially for populations with low-income.^33^

The results highlighted that a majority of the improperly masked or unmasked were male or middle age adults in the approximate age range of 25 to 44 years (**Figs 4 and 5**). Finding a high prevalence of incorrect mask use among males coincides with previous research findings. A number of studies report that females are more likely than males to report wearing masks or wearing masks appropriately.^15,29,34^ One study suggests a connection between masculinity and mask-wearing behaviors^30^ and other research suggests that the tendency to appropriately wear a mask is connected to one’s caregiving responsibilities.^35^ Some have argued that masking-wearing as a public health practice may evolve like the utilization of seatbelts did beginning in the mid-1900s in the United States. If so, identifying the history of successful and unsuccessful public health practice campaigns will further public marketing for masking. For example, researchers have reported that young men are the most unlikely to follow seatbelt regulations,^36-38^ and others have suggested that seatbelt use is connected with one’s perceived risk.^39^ Even other public health prevention tools, such as condom use, has been connected with one’s tendency to take risk or engage in impulsive behaviors.^40^ This evidence may suggest mask-wearing behaviors may face similar challenges.

The geographical and cultural context in which this study observed mask-wearing behaviors warrants attention. One survey identified the political and polarizing nature of mask-wearing among United States counties.^41^ Louisville is an urban area within a predominately rural state with midwestern and southern history. Mask-wearing behaviors may differ between rural and urban settings as well.^29^ Louisville voted for democratic representatives in the House of Representatives eight out of the past ten election cycles.^42^ In 2018, Kentucky elected a democratic governor after four years with a republican governor^43^ which speaks to the dynamic political and cultural context of the state. Thus, a wider sampling of Kentucky as a whole state would provide a more representative snapshot of these behaviors to the geographical and political variation in the state.

### Limitations

Only businesses classified as Retail Trade by SIC were observed in this study. Retail trade establishments make about 15% of all business establishments in the U.S., about 14% in Louisville.^22,23^ Therefore, the result of this study is not representative of mask-wearing behavior in indoor public areas of non-retail trade businesses. Even among the retail trade, two major classifications − namely, (1) automotive dealers and gasoline service stations and (2) eating and drinking places, constituting about 5% of all businesses in Louisville-were excluded.^23^ Observing eating and drinking places is especially important to understand the dynamics of the spread of respiratory infectious disease, as they are environments where masks are taken off, at least occasionally, and have been linked to an increase in COVID-19 cases.^44^

The representativeness of the observed sample of PAs was not as complete as the study strategically planned, though the error was still small. The median representativeness error (defined as the difference of the share of observed PA of a specific size from a district in total observed PAs of that size from the population share of the district) in the observed sample was 3.92%. Approximately a half of the error could be attributed to the sample selection mechanism that resulted in a median representativeness error of 1.95%. The rest could be attributed to the implementation challenges. For example, some of the selected PAs were either non-operational or could not be visited by surveyors. This happened more often in small PAs (square feet less than 1499), especially in the West Center, North Center, and South East Districts.

This study’s results for sex and age-range of the unmasked and incorrectly masked need careful interpretation. For example, the majority of the unmasked and incorrectly masked visitors of the observed PAs were male and middle age adults. However, one cannot be certain that males and middle age adults exhibit the worst mask-wearing behavior without confirming the sex and age of the masked and correctly mask visitors as well. The later pieces of information were not directly collected from the visitors and staff in this study. Nonetheless, the cross-district variation in mask use by the demographic characteristics is more informative than the overall prevalence.

## Conclusion

The findings from this observational study showed that the incorrectly masked and unmasked visitors were more commonly observed in small public areas (Square footage < 1,500). In contrast, the incorrectly masked and unmasked staff were more commonly observed in medium-small public areas (Square footage = 1500 - 2500). Both incorrectly masked visitors and staff were most commonly observed in food and grocery stores than other retail stores. Among the observed visitors and the staff, middle age adults made up the highest proportion of unmasked and incorrectly masked. Despite mask mandates in place, we observed a small proportion of visitors (4%) and staff (8%) that did not wear a facial mask or wore them incorrectly (14% of visitors and 13% of staff). There is a continued need to improve awareness of the effectiveness of appropriate facial mask use, financial resources to provide masks, particularly in low-income areas, and education on correct ways to wear a mask.

## Author Contributions

1. **Seyed M. Karimi:** Conceptualization, Methodology, Fieldwork organization, Surveyor, Data curation, Formal analysis, Supervision, Writing the original draft, Reviewing and editing
2. **Sonali S. Salunkhe:** Conceptualization, Methodology, Fieldwork organization, Surveyor, Data curation, Formal analysis, Supervision, Writing the original draft, Reviewing and editing
3. **Kelsey White:** Conceptualization, Methodology, Fieldwork organization, Surveyor, Data curation, Formal analysis, Supervision, Writing the original draft, Reviewing and editing
4. **Bert B. Little**: Conceptualization, Methodology, Supervision, Reviewing and editing
5. **W. Paul McKinney:** Conceptualization, Methodology, Supervision, Reviewing and editing
6. **Riten Mitra:** Conceptualization, Methodology, Supervision, Reviewing and editing
7. **YuTing Chen:** Methodology, Data Curation
8. **Emily R. Adkins:** Surveyor
9. **Julia A. Barclay:** Surveyor
10. **Emmanuel Ezekekwu:** Surveyor
11. **Caleb X. He:** Surveyor
12. **Dylan M. Hurst:** Surveyor
13. **Martha M. Popescu:** Surveyor
14. **Devin N Swinney:** Surveyor
15. **David Johnson:** Organization of the survey’s logistics at the University of Louisville
16. **Rebecca Hollenbach**: Organization of the survey’s logistics at the LMPHW
17. **Sarah Moyer:** Organization of the survey’s logistics at the LMPHW
18. **Natalie C. DuPré:** Conceptualization, Methodology, Supervision, Reviewing and editing

## Acknowledgments

Seyed Karimi, Ph.D., the corresponding author of this manuscript, is responsible for (is the guarantor of) the content of the manuscript, including the data and analysis. The Louisville Metro Department of Public Health and Wellness (LMPHW) funded this study through the Coronavirus Aid, Relief, and Economic Security Act (the CARES Act). The Commonwealth Institute of Kentucky (CIK) provided access to Qualtrics to design this study’s online questionnaire.

